# Foot-and-mouth disease virus localisation on follicular dendritic cells and sustained induction of neutralising antibodies is dependent on binding to complement receptors (CR2/CR1)

**DOI:** 10.1101/2021.09.08.459380

**Authors:** Lucy Gordon, Neil Mabbott, Joanna Wells, Liudmila Kulik, Nick Juleff, Bryan Charleston, Eva Perez-Martin

## Abstract

Previous studies have shown after the resolution of acute infection and viraemia, foot-and-mouth disease virus (FMDV) capsid proteins and/or genome are localised in the light zone of germinal centres of lymphoid tissue in cattle and African buffalo. The pattern of staining for FMDV proteins was consistent with the virus binding to follicular dendritic cells (FDCs). We have now demonstrated a similar pattern of FMDV protein staining in mouse spleens after acute infection and showed FMDV proteins are colocalised with FDCs. Blocking antigen binding to complement receptor type 2 and 1 (CR2/CR1) prior to infection with FMDV significantly reduced the detection of viral proteins on FDCs and FMDV genomic RNA in spleen samples. Blocking the receptors prior to infection also significantly reduced neutralising antibody titres. Therefore, the binding of FMDV to FDCs and sustained induction of neutralising antibody responses are dependent on FMDV binding to CR2/CR1 in mice.

**Author Summary:** Foot and mouth disease virus causes a highly contagious acute vesicular disease, resulting in more than 50% of cattle, regardless of vaccination status, and almost 100% of African buffalo becoming persistently infected for long periods (months) of time. Yet, the mechanisms associated with establishment of persistent infections are still poorly understood. Infected animals are characterised by the presence of long-lived neutralising antibody titres, which contrast with the short-lived response induced by vaccination. We have used a mouse model to understand how foot and mouth disease virus is trapped and retained in the spleen for up to 28 days post infection and how the absence of antigen in the germinal centre prevents a sustainable neutralising antibody response, in the mouse. Our results highlight the importance of targeting antigen to FDCs to stimulate potent neutralising antibody responses after vaccination.

## Introduction

One of the features of foot-and-mouth disease virus (FMDV) infection, which has a major impact on the control and eradication of foot-and-mouth disease (FMD), is the existence of the “carrier state” (1, 2). A carrier of FMDV is defined as an animal from which live virus can be recovered from the nasopharynx after 28 days following infection, which frequently occurs in ruminants after acute infection (3). Only ruminants are considered FMDV carriers, and among them, around 70% of infected African buffalo become carriers after acute infection and can carry FMDV for up to 5 years or more, which is why African buffalo are considered the primary reservoir of FMDV in Africa (4–6). Over 50% of cattle exposed to FMDV become carriers (4, 5, 7), and although current vaccines prevent clinical disease, they do not prevent primary infection in the nasopharynx, therefore vaccinated animals can still become persistently infected carriers of FMDV (8).

FMDV infection in ruminants elicits the production of specific serum neutralising antibody titres which can provide protection for years (6, 9). T cell depletion studies in cattle identified that CD4^+^ T-cell-independent antibody responses are required for resolution of clinical FMD in cattle (10). Similarly, FMDV vaccines induce predominantly CD4^+^ T-independent antibody responses that are enhanced by T cell activation (11). Current inactivated FMD vaccines generally offer only a short-lived immune response in the host, due to the inability to induce FMDV-specific memory B cells. Neither infection nor vaccination induces a significant number of circulating memory B cells, despite a key difference of longer duration of immunity post-infection compared to post-vaccination (12).

Antigen retention on stromal follicular dendritic cells (FDCs) has been shown to maintain humoral immune responses by retaining complement-coated immune complexes (ICs) on their surface for long periods of time via complement receptors (CR2/CR1) and/or antibody Fc receptors (13–15). Juleff et al. first hypothesised that, unlike vaccination, upon natural infection FMDV binds to and is retained by FDCs in the form of immune-complexed FMDV particles, resulting in prolonged stimulation of short-lived plasma cells, which maintains high levels of neutralising antibodies (16). FDCs are specialised immune cells of stromal origin found in the spleen, lymph nodes (LNs) and other lymphoid tissue including tonsil and mucosal surfaces, within B cell follicles in the light zones of germinal centres (GCs) (17). They are necessary for GC formation, lymphoid follicle organisation and promoting B cell proliferation, survival and differentiation (18). FDCs present ICs to both naïve and GC B cells; therefore together with B cells, FDCs are crucial for an effective humoral immune response (19). The longevity of FDCs and their ability to trap and retain antigens has also been exploited by certain pathogens. FDCs represent a major extracellular reservoir for a number of viruses and other pathogens including, but not limited to, human immunodeficiency virus (HIV), vesicular stomatitis virus (VSV), bovine viral diarrhoea virus (BVDV) and prions (20–23).

With regards to FMDV infection in cattle, it was demonstrated that the virus can persist in association with the light zone FDC network of GCs in lymphoid tissues of the head and neck (16). The data provided potential insight into both the mechanisms of viral persistence and the long-lasting antibody responses seen upon natural infection. An alternative study has described the site of FMDV persistence as pharyngeal epithelial cells in both vaccinated and non-vaccinated persistently infected cattle within the mucosa-associated lymphoid tissue, interestingly associated with CR2^+^ sub-epithelial lymphoid follicles (24). We have also found that in buffalo persistently infected with the Southern African Territories (SAT) FMDV serotypes SAT-1, SAT-2 and SAT-3, quantities of FMDV RNA were significantly higher in GCs in lymphoid tissue compared to epithelium samples, which again warranted further investigation into the possibility of virus-persistence in association with FDCs (25).

Data from experiments in mice have been fundamental in demonstrating the complement receptor-mediated retention of certain pathogens on FDCs (26, 27). For example, Ho et al. were able to demonstrate the binding of HIV to lymph node FDCs by using a rat monoclonal antibody (mAb) 7G6 to block CR2, which in turn prevented binding and retention of virions (28). This observation was confirmed with the use of CR2/CR1- deficient (*Cr2^−/−^*) mice, whereby no virus could be detected on FDCs (28).

Using a mouse model to FMDV persistence, our previous data suggested that splenic FDCs were able to trap and maintain FMDV in the GC light zone for up to 63 dpi (29). The main aim of our study was to identify the receptor(s) involved in the maintenance of FMDV antigen (Ag) within the GC, and whether retention of Ag impacted the generation and maintenance of neutralising antibodies to FMDV in mice. We show that the blocking of CR2/CR1 on FDCs prevented binding and retention of FMDV, strongly suggesting this interaction is mediated by FMDV binding to CR2/CR1. Further investigation using super-resolution microscopy showed significant co-localisation of FMDV Ag with CR2/CR1^+^ FDCs in the spleen. Moreover, blocking of CR2/CR1, and consequently absence of FMDV Ag on FDCs, resulted in the significant reduction of neutralising antibody responses to FMDV. A key function of FDCs in the GC reaction is the presentation of antigen, in the form of ICs, to B cells, driving affinity maturation. Blocking CR2/CR1 resulted in antibodies with a reduced capacity to neutralise virus and lower binding affinity to FMD virus-like particles (VLPs) compared to control animals. Until now, CR2/CR1 were not known to bind and maintain FMDV Ag on FDCs, resulting in the production of high avidity, neutralising antibodies; therefore, knowledge of this interaction could enable a targeted approach to vaccine design, incorporating the binding of complement-coated FMDV-ICs on FDCs via CR2/CR1 to increase duration of immunity post-vaccination.

## Results

### Monoclonal antibody (mAb) 4B2 binds CR2/CR1^+^ in Balb/C mice and does not affect the proportion of immune cells in the spleen

In order to study antigen retention, a mouse anti-CR2/CR1 mAb 4B2 was used, which had been shown to block CR2/CR1 for up to 6 weeks *in vivo* in C57 Black mice, thus an excellent reagent for studying long term persistence of FMDV on FDCs in mice (30). First, mice were injected with mAb 4B2, or IgG1 as an isotype control, and effects on splenocytes determined by flow cytometry at intervals afterwards up to 35 days post injection. We chose 3 anti-CR mAbs to test the blocking ability of mAb 4B2 up to 35 days post injection. The most notable reduction was the binding of mAb 7G6 to splenocytes from 2 days post inoculation (Fig 1A). This mAb binds a similar and overlapping epitope on CR1 and CR2 as mAb 4B2 as described previously (30).

**Fig 1.**
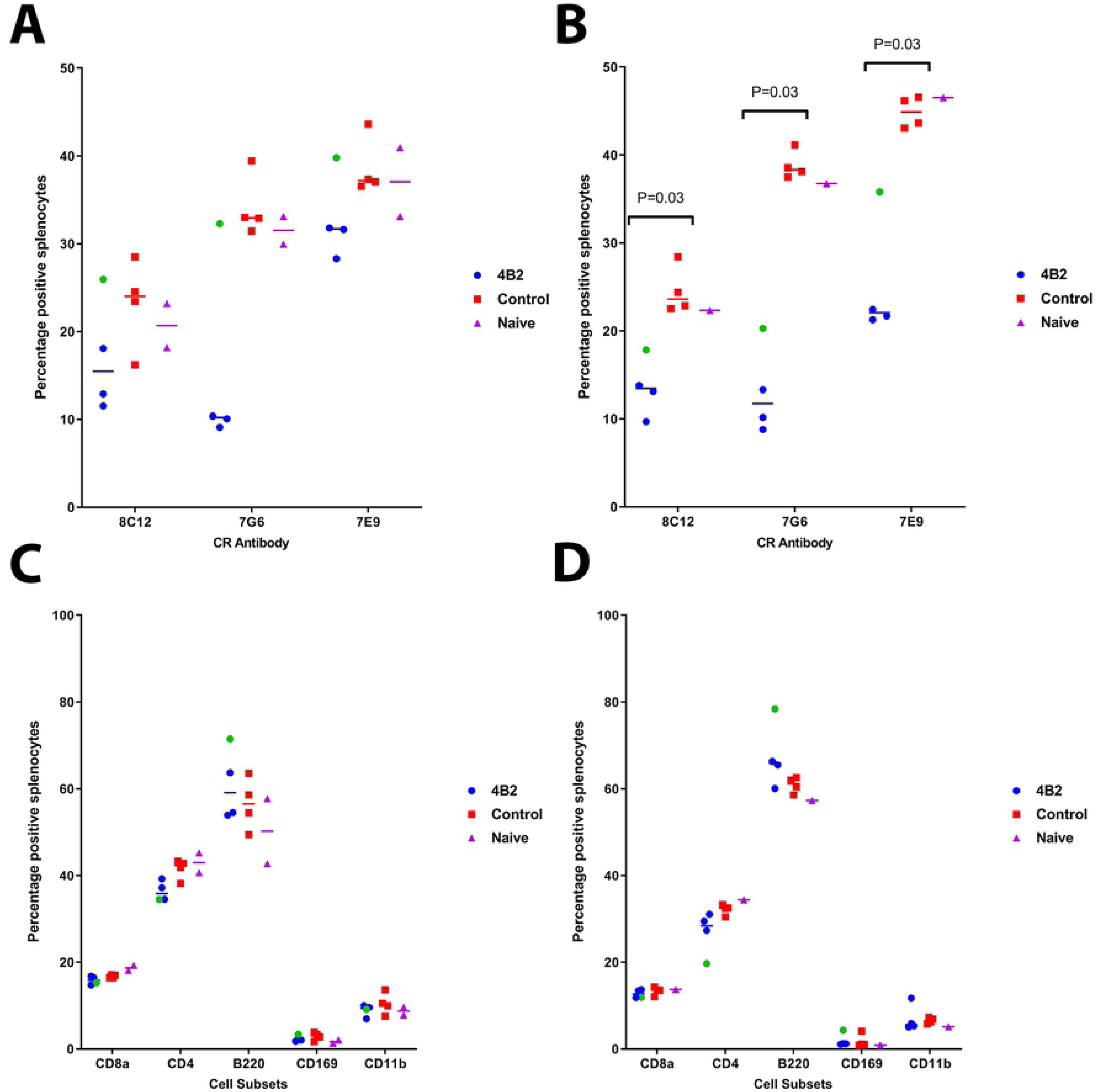
Flow cytometric analysis of splenocytes from mice treated with 4B2 mAb or control IgG, comparing cell subsets and availability of CR.

Flow cytometry was used to identify availability of complement receptors in mice after treatment with mAb 4B2 and the percentage of cell subsets, compared to control mice treated with IgG1. Spleen samples were taken at (**A**, **C**) early time points and (**B**, **D**) late time points from mice treated with 4B2 or IgG1 and naïve animals. At the early time points (**A**) there is a trend whereby mice treated with mAb 4B2 show a smaller number of positive cells to the CR antibodies, compared to the IgG1 or control groups, although this is not significant; 8C12 p = 0.312; 7G6 p = 0.061; 7E9 p = 0.194. By the late time points (**B**) mice treated with 4B2 had significantly reduced binding of the 3 anti-CR antibodies (p = 0.03) to their cells compared to the control mice. The percentage of the different splenic cell subsets CD8 and CD4 T cells, B cells (B220), macrophages (CD169) and dendritic cells (CD11b) (**C**-**D**) remained unchanged after treatment with 4B2 when analysed from both early and late time points after antibody treatment. Naïve animals were used as controls as they were untreated. Splenocytes analysed from 1 mouse at the early time point and 1 mouse from late time point are highlighted in green to show the mice where the 4B2 antibody treatment didn’t seem to work and their values throughout the flow data. These mice were included in all statistics. *p values are 0.03 using the non-parametric Mann-Whitney U test to compare the medians of the two treatment groups.

Up to 35 days post inoculation mAb 4B2 is still capable of blocking CR, with a significant reduction of anti-CR2/CR1 mAbs binding to splenocytes, as demonstrated not only by mAb 7G6, but mAbs 8C12, which is monospecific to CR1, and 7E9, which binds a different epitope on CR2/CR1 (Fig 1B). This is in alignment with previous data, whereby the blocking effect is not purely due to steric inhibition, but induces a substantial decrease in the expression level of receptors when mAb 4B2 is used *in vivo* (30).

Treatment with mAb 4B2 did not affect the abundance of CD8a cytotoxic T cells, CD4 T helper cells, B cells (B220+ cells), marginal zone macrophages (CD169) and monocytes (CD11b), at early or late timepoints (Figs 1C and 1D respectively). Importantly, immunohistochemistry (IHC) analysis showed the presence of CD21/CD35+ FDCs in the spleens of mice treated with mAb 4B2 (Figs 2 and 4). These data are consistent with data from Kulik et al. that also reported that i*n vivo* injection of mAb 4B2 does not induce the death of immune cells including FDCs, but leads to substantial blocking of binding of other mAb to CR2 and CR1 (30).

**Fig 2.**
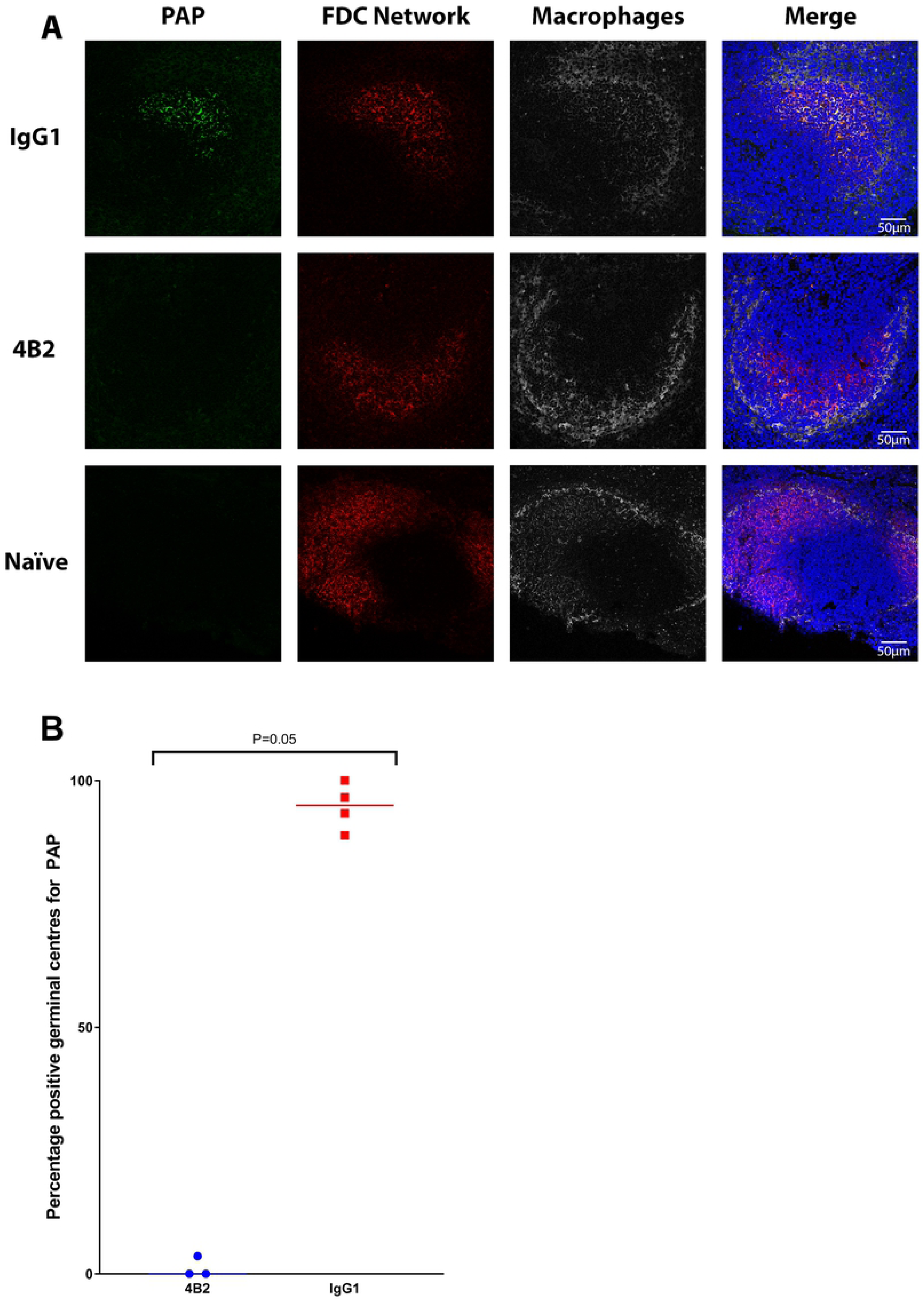
Effect of pre-treatment with 4B2 preventing the binding of PAP on the FDC network within the GC in spleen.

### Reduced immune complex trapping by FDC in the spleens of mice treated with mAb 4B2

We next investigated the effects of *in vivo* mAb 4B2 treatment on the ability of FDC to trap immune complexes. Mice were injected with mAb 4B2 (or an IgG1 isotype control) and 1 day later injected with pre-formed peroxidase-anti-peroxidase (PAP) containing ICs which can bind to FDCs *in vivo* via CR2/CR1 (31). Spleen sections were analysed by confocal microscopy 1 day later (Fig 2). The presence of CR2/CR1-expressing FDC was detected using mAb 7E9. In control-treated mice, PAP-ICs were consistently detected in association with FDC in 95% of the splenic light zone GCs examined, with just 5% of GCs negative for PAP. In contrast, in the spleens of mice treated with mAb 4B2, PAP-ICs were detected in fewer than 2% of the GCs examined, similar to the background levels observed in naïve mice (Table 1). This data demonstrates that pre-treatment of mice with mAb 4B2 effectively blocks the retention of ICs by splenic FDCs *in vivo*.

**Table 1.**
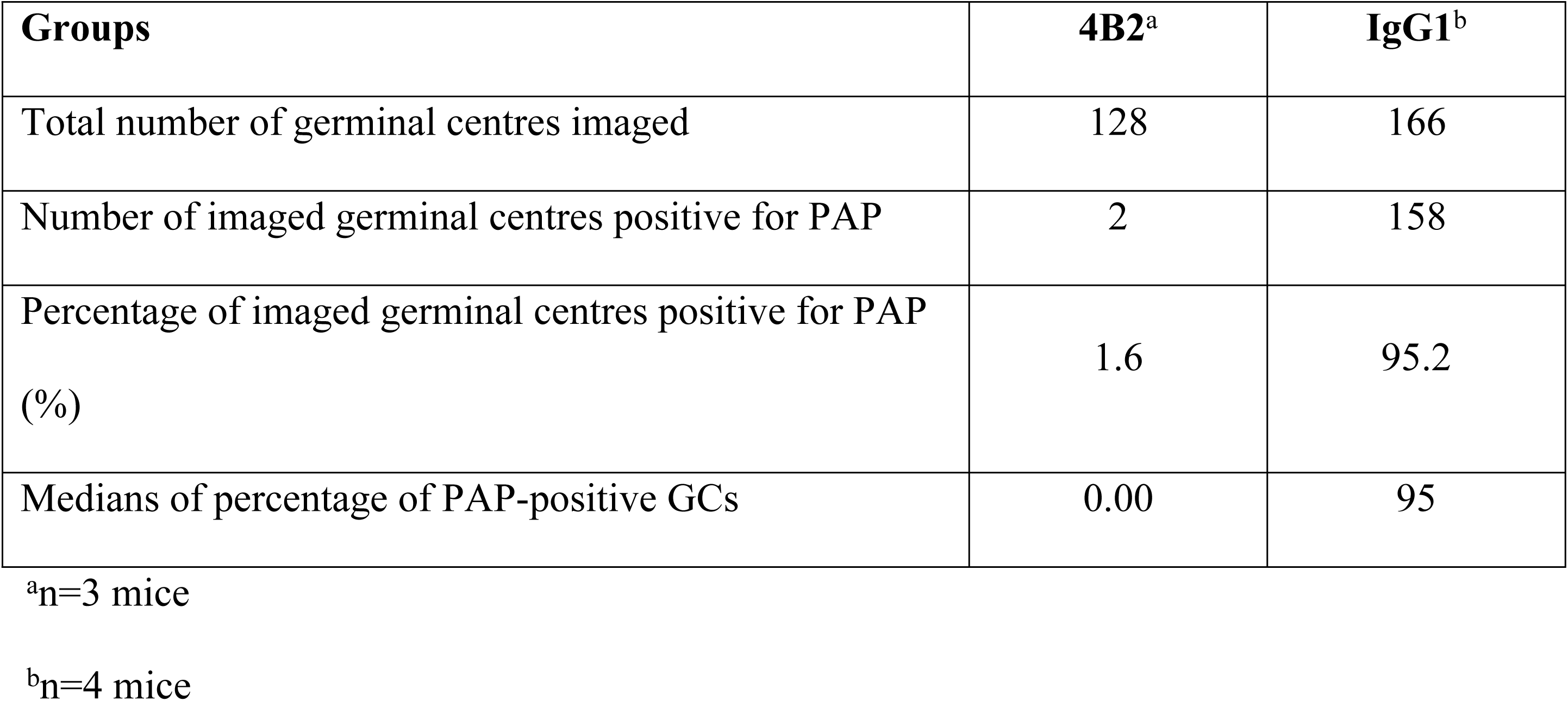
Immunofluorescence examination of GCs in mice spleens treated with CR block (4B2) or an isotype control (IgG1) 1 day before PAP immunisation.

BALB/c mice were treated with 500 μg of 4B2 (n=3) or IgG1 (n=4) control 24hr before immunisation intravenously with peroxidase anti-peroxidase (PAP). Naïve mice were untreated. Spleen samples were collected in O.C.T from mice culled 1-day post inoculation with PAP. **A**) Cryosections were analysed via confocal microscopy for the presence of PAP within the light zone of the GCs associated to the FDC network. Confocal microscopy images are arranged in rows and columns according to the treatment and the staining. **B**) Mice from the 4B2 treatment group had significantly less PAP bound to FDCs compared the control group, p value of 0.05 using the non-parametric Mann-Whitney U test to compare the medians of the two treatment groups.

**PAP panels**: show PAP labelled green, detected with anti-rabbit 488. PAP was detected in the light zone GC associated to the FDCs in IgG1 control mice, but not in 4B2 treated mice. Absence of signal to PAP in naïve mouse spleen.

**FDC Network**: show light zone FDCs labelled red with Alexa Fluor 594-conjugated anti-mouse CD21/CD35 (CR2/CR1) antibody, clone 7E9; FDC clusters were detected in all groups.

**CD169 panels**: show marginal zone macrophages surrounding the light zone GC labelled grey with conjugated mAb CD169-APC.

**Merged images**: show deposition of PAP within the light zone FDCs network (yellow, co-localisation) of the IgG1 control mice but not in 4B2 treated mice or naïve mice. Nuclei stained blue (DAPI). Scale bars = 50 μm.

### CR2/CR1-blockade enhances the viraemia during FMDV infection

Next, we determined the effects of mAb 4B2-mediated CR2/CR1-blockade on the viraemia during FMDV infection. Mice were treated with mAb 4B2, or IgG1 as a control, and 1 day later injected with FMDV. By two days after infection a statistically significant, 10-fold increase in the viraemia in sera was detected in mAb 4B2-treated mice compared to control-treated mice (Fig 3A). Viral RNA quantification corroborated the plaque assay results, whereby mAb 4B2-treated mice showed a statistically significant, 10-fold increase of viral genome in the serum compared to the control-treated mice (Fig 3B). By 7 dpi the viraemia was cleared in both groups and no detectable virus was detected by plaque assay or qPCR. Naïve mice (n=3) were used as negative controls and were negative for both the plaque assay and qPCR (data not shown). These data show that blockade of CR2/CR1 resulted in a higher titre of virus in sera post-infection with FMDV in mice.

**Fig 3.**
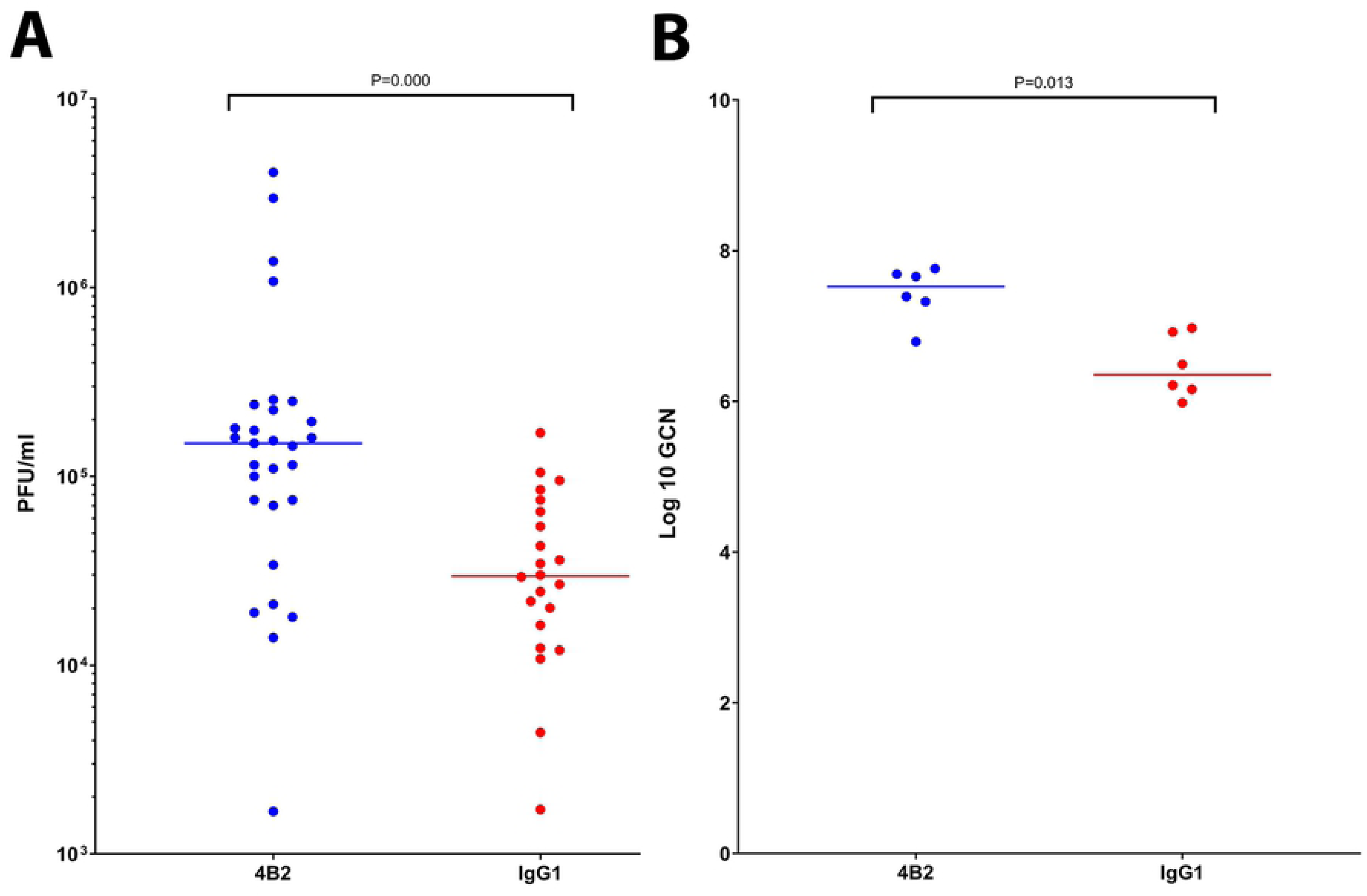
Viraemia in 4B2 treated and control-treated mice in response to FMDV infection.

The presence of viraemia in serum samples from mice treated with 4B2 or IgG1 was investigated by (**A**) plaque assay and (**B**) qRT-PCR. Serum samples were collected from 4B2 and IgG1 treated mice at 2 and 7 dpi. The quantity of virus in the mouse serum at 2 dpi is expressed as (**A**) the number of plaque forming units per 1 ml serum and (**B**) log 10 genome copy number. Each blot represents an individual animal, and the line represents the median values. Naïve mice at 2 dpi and all serum samples harvested from 7 dpi were negative for viraemia (data not shown). P values of <0.05 using the non-parametric Mann-Whitney U test to compare the medians of the two treatment groups.

### CR2/CR1-blockade reduces the trapping and persistence of FDMV antigen in the spleen

Next, we determined whether CR2/CR1 blockade similarly impeded the trapping and persistence of FMDV in the spleen. Mice were treated with mAb 4B2 or IgG1, 1 day later injected with FMDV, and spleens (n=8/group) collected at weekly intervals afterwards.

Spleens from naïve mice were used as controls. The location of FMDV and FDC networks in the spleens was determined by immunofluorescence confocal microscopy (Fig 4 A-D). We used mAb 7E9 to detect FDC since the treatment of mice with mAb 4B2 does not completely block the binding of mAb 7E9 to CR2/CR1 (Figs 1-2). The total number of FDC networks positive or negative for FMDV is represented in Table 2.

**Fig 4.**
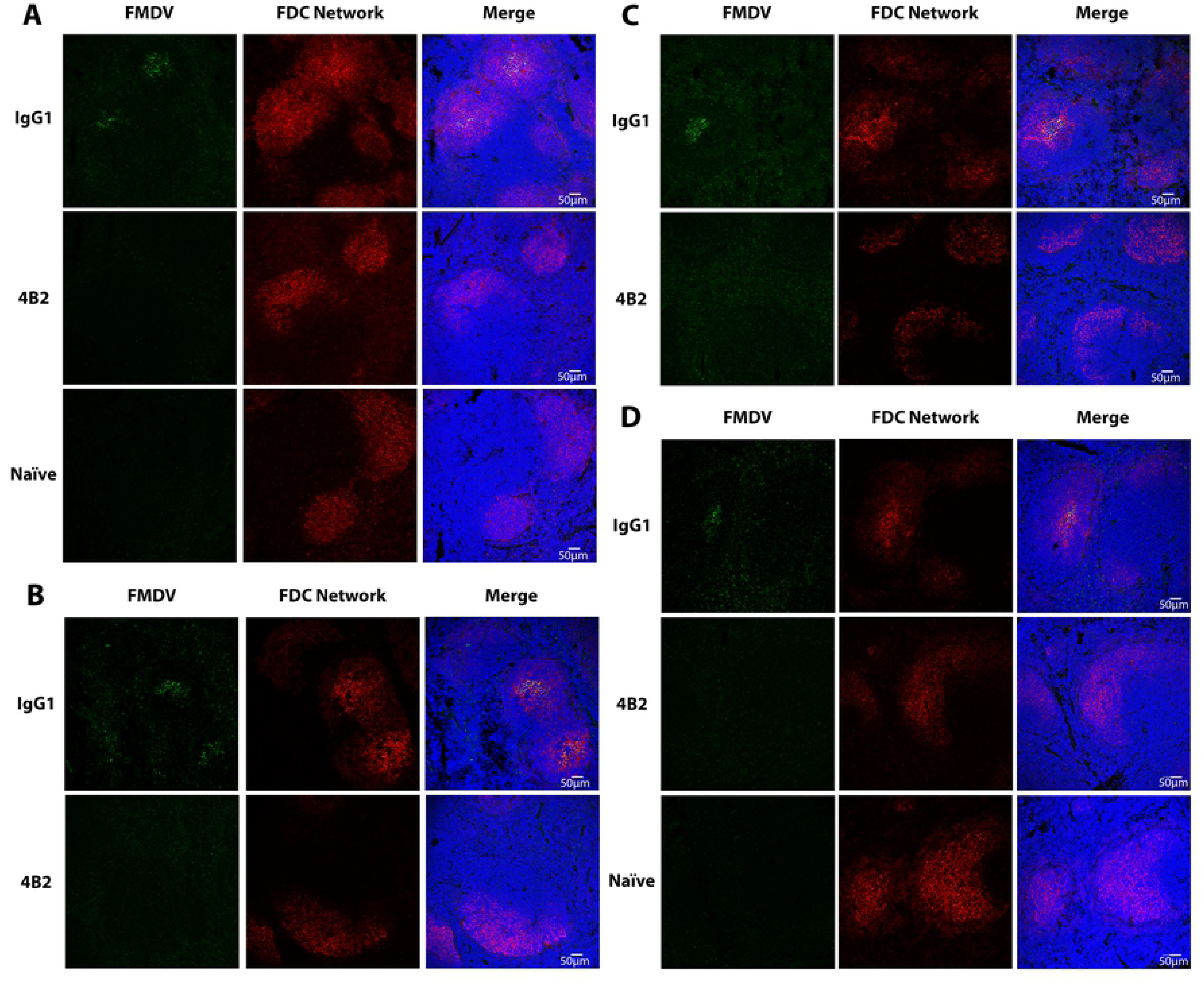
Effect of pre-treatment with 4B2 preventing the binding of FMDV on the FDC network within the GC in mouse spleen.

**Table 2.**
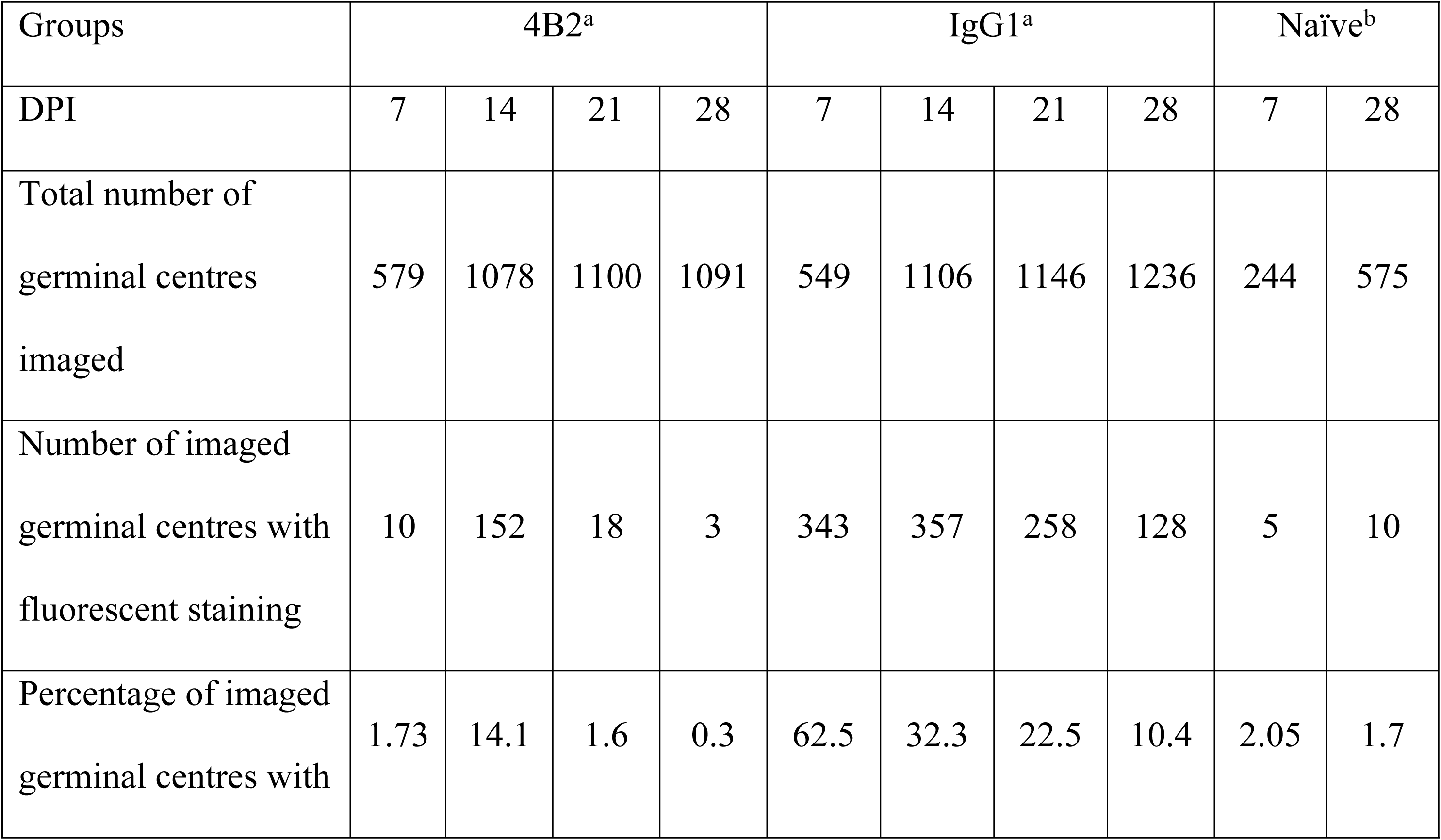
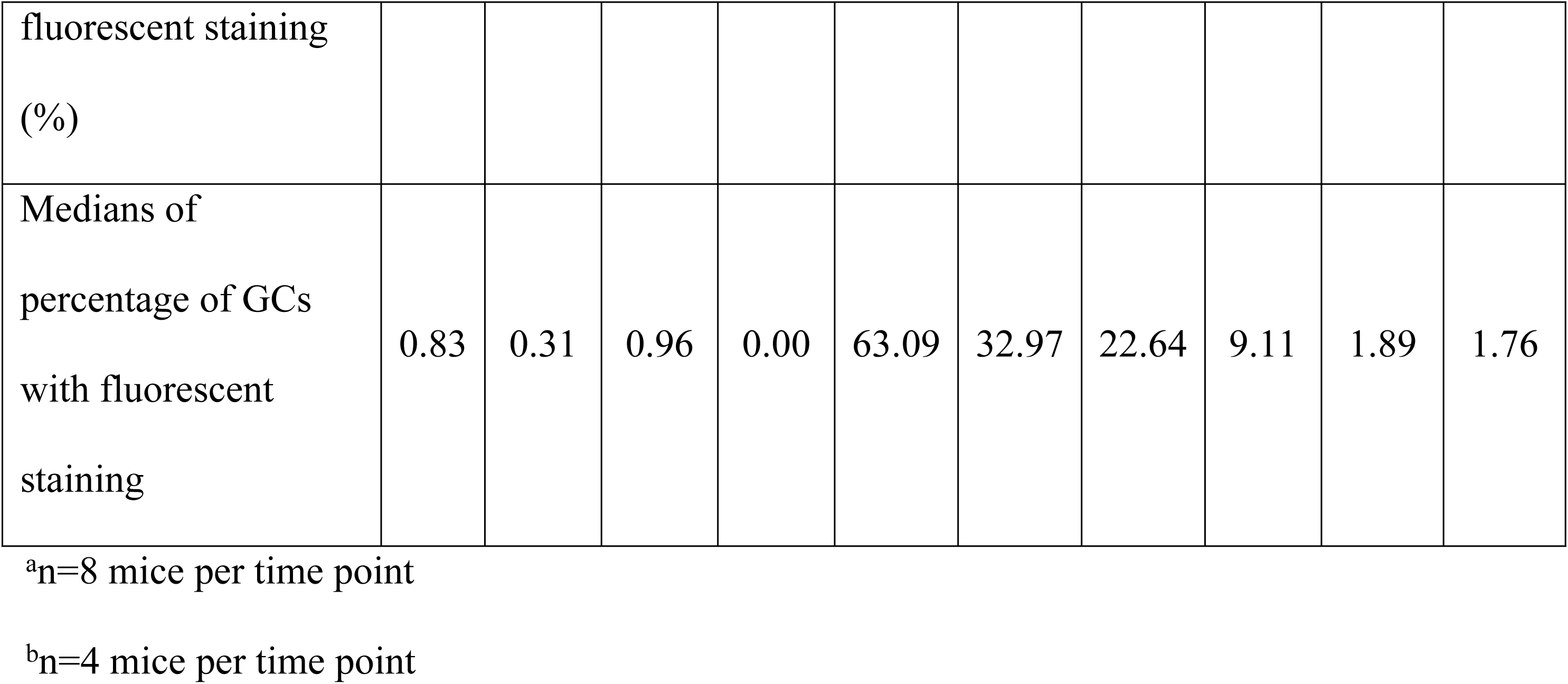
Immunofluorescence examination of mouse spleen treated with CR block or control IgG1 for FMDV in light zone GCs at 7, 14, 21 and 28 days post infection; and naïve mice at 7 and 28 dpi.

| Groups | 4B2<sup>a</sup> |  |  |  | IgG1<sup>a</sup> |  |  |  | Naïve<sup>b</sup> |  |
| --- | --- | --- | --- | --- | --- | --- | --- | --- | --- | --- |
| DPI | 7 | 14 | 21 | 28 | 7 | 14 | 21 | 28 | 7 | 28 |
| Total number of<br>germinal centres<br>imaged | 579 | 1078 | 1100 | 1091 | 549 | 1106 | 1146 | 1236 | 244 | 575 |
| Number of imaged<br>germinal centres with<br>fluorescent staining | 10 | 152 | 18 | 3 | 343 | 357 | 258 | 128 | 5 | 10 |
| Percentage of imaged<br>germinal centres with | 1.73 | 14.1 | 1.6 | 0.3 | 62.5 | 32.3 | 22.5 | 10.4 | 2.05 | 1.7 |
| fluorescent staining<br>(%) |  |  |  |  |  |  |  |  |  |  |
| Medians of<br>percentage of GCs<br>with fluorescent<br>staining | 0.83 | 0.31 | 0.96 | 0.00 | 63.09 | 32.97 | 22.64 | 9.11 | 1.89 | 1.76 |

<sup>a</sup>n=8 mice per time point
<sup>b</sup>n=4 mice per time point

Confocal microscopy images are arranged in rows and columns according to the treatment, day post infection and the staining. BALB/c mice treated with 500 μg of either 4B2 mAb or IgG1 control mAb on day -1 before challenge with FMDV. FMDV-infected mouse spleen samples were collected at (**A**) 7, (**B**) 14, (**C**) 21 and (**D**) 28 days post infection from IgG control mice and 4B2 mice (n=8 per group per timepoint). Naïve mouse spleens were taken at 7 dpi (n=4) and 28 dpi (n=4).

**FMDV panels**: show FMDV protein labelled green with biotinylated llama single domain anti-FMDV 12S antibody VHH-M3 and streptavidin Alexa-Fluor-488. FMDV was detected in the IgG1 control group at all timepoints. FMDV was not detected in the spleens of the 4B2 treated group at any of the time points, with the exception of four mice, three harvested at day 14 and one at day 21. There was an absence of signal for FMDV in naïve mouse spleen at all time points.

**FDC Network**: show FDCs in the light zone GC labelled red with Alexa Fluor 594-conjugated anti-mouse CD21/CD35 (CR2/CR1) antibody, clone 7E9; FDC clusters were detected at all time points in control, 4B2 treated and naïve mice.

**Merged images**: shows deposition of FMDV within the light zone FDC network of IgG1 control mice (yellow – colocalization); but absence of FMDV within the light zone FDC network of 4B2 treated mice and naïve mice (red). Nuclei stained blue (DAPI). Scale bars = 50 μm.

In IgG1 isotype control treated mice, FMDV-Ag was detected in the majority of FDC networks by 7 dpi. Although number of FMDV-Ag-positive FDC networks gradually declined as the infection progressed, FMDV-Ag was detectable in association with approximately 10% of the FDC networks at 28 dpi (Fig 5A). In contrast, no association of FMDV-Ag with FDCs above background levels was detected in the spleens of mAb 4B2 treated mice, with the exception of 4 mice, suggesting that CR2/CR1-blockade had prevented the trapping and retention of FMDV on FDC. The treatment may not have been completely effective in the 4 anomalous mice, 3 killed at 14 dpi and 1 killed at 21 dpi (represented as green dots in Figs 5-9).

**Fig 5.**
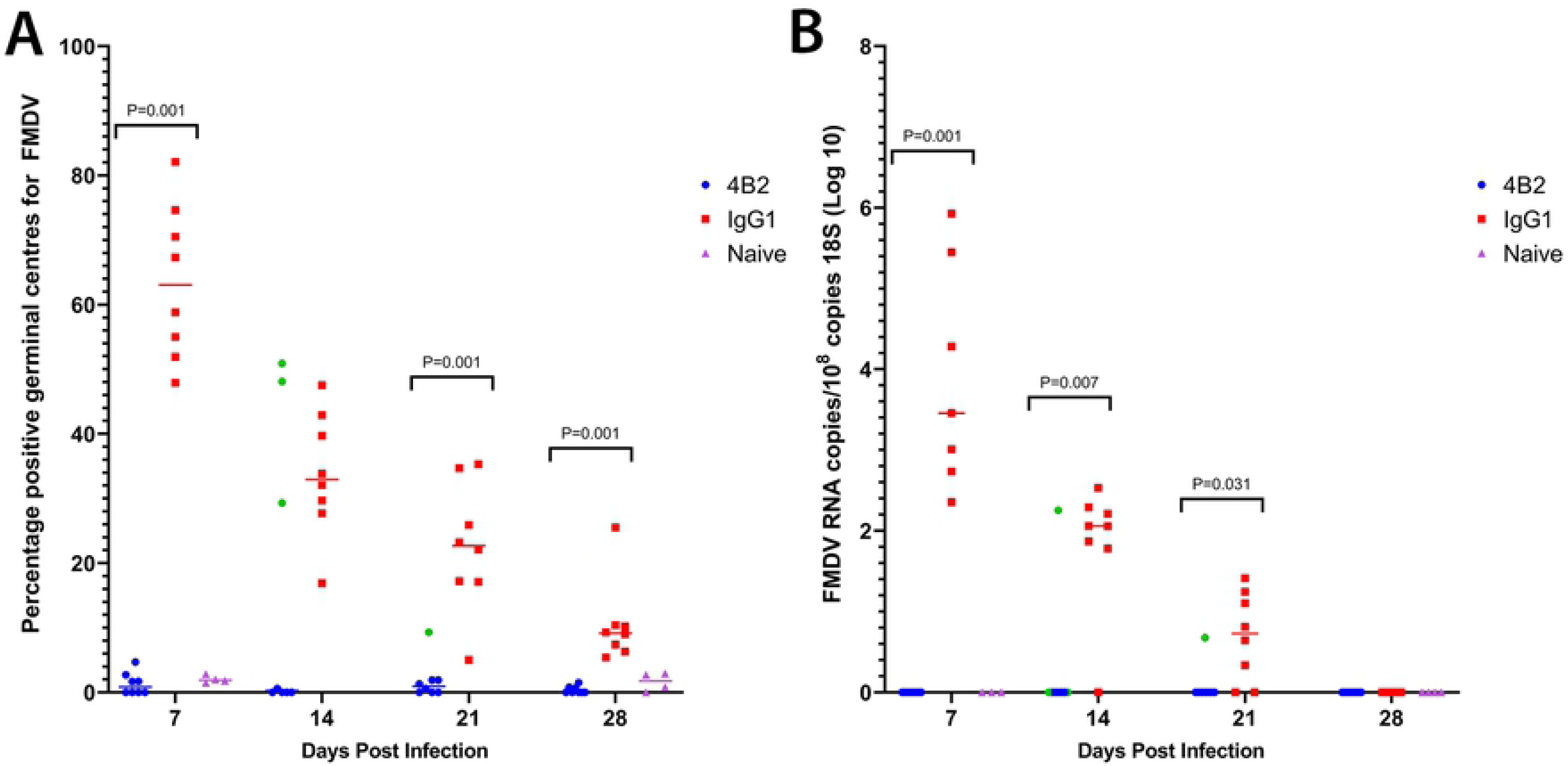
Quantification of FMDV antigen and RNA in spleens from 4B2-treated and isotype control treated mice detected by confocal microscopy and RT-qPCR.

Spleen samples were collected from BALB/c mice at 7, 14, 21 and 28 days post infection (n=8/group/timepoint) following treatment with either 4B2 mAb or IgG1 isotype control mAb one day prior to IP challenge with 10^6.2^ TCID_50_ of FMDV/O/UKG/34/2001.

**A)** Microscopic analysis of spleen sections demonstrated that mice treated with 4B2 had significantly less FMDV in their GCs, with a P value of ≤0.001 from 7, 21 and 28 post infection. GCs were visualised and imaged using mAb 7E9, an anti-CR2/CR1 antibody. The GCs were counted and the percentage which were positive for FMDV, detected using a biotinylated llama single domain anti-FMDV 12S antibody VHH-M3, was calculated.

**B)** The samples were analysed by RT-qPCR for the presence of FMDV RNA and the results are expressed as copies per 10^8^ copies of 18S rRNA. Each point represents an individual animal and the line represents the median values. CT values ≥ 35 for 3D FMDV were deemed negative and recorded as 0. Naïve mice (n=4) were tested at 7 and 28 dpi as negative controls. Using the non-parametric Mann-Whitney *U* test to compare the medians of the two groups, at 7 dpi p value of 0.001; 14 dpi p value of 0.007; 21 dpi p value of 0.031.

Comparison of the presence of viral RNA similarly revealed that CR2/CR1-blockade had prevented the accumulation and persistence of FMDV in the spleen. While high levels of viral RNA were detected in the spleens of control-treated mice until 21 dpi, the levels in 4B2-treated mice were below the detection limit (Fig 5B). However, although FMDV-Ag was detectable in association with FDCs in the spleens of control-treated mice by 28 dpi, the levels of viral RNA in whole spleen samples were below the detection limit in all groups at this time. Thus, these data show that trapping and persistence of FMDV Ag is dependent on FMDV binding to FDCs via CR2/CR1.

### Co-localisation of CR2/CR1 with FMDV

Localisation of FMDV was consistently found in murine spleens within the FDC networks. Further investigation using stimulated emission depletion (STED) microscopy for super-resolution images confirmed that FMDV proteins were predominantly co-localised with CR2/CR1 on FDCs (Fig 6). ImageJ software was used to confirm that the distribution of the CR2/CR1- and FMDV-Ag-associated fluorochromes were preferentially co-localised, compared to that predicted by the null hypothesis that each of these were randomly and independently distributed (31, 32). This analysis confirmed a highly significant and preferential association of the FMDV-Ag with CR2/CR1 on FDCs when compared to the null hypothesis that the pixels were randomly distributed (Fig 6B).

**Fig 6.**
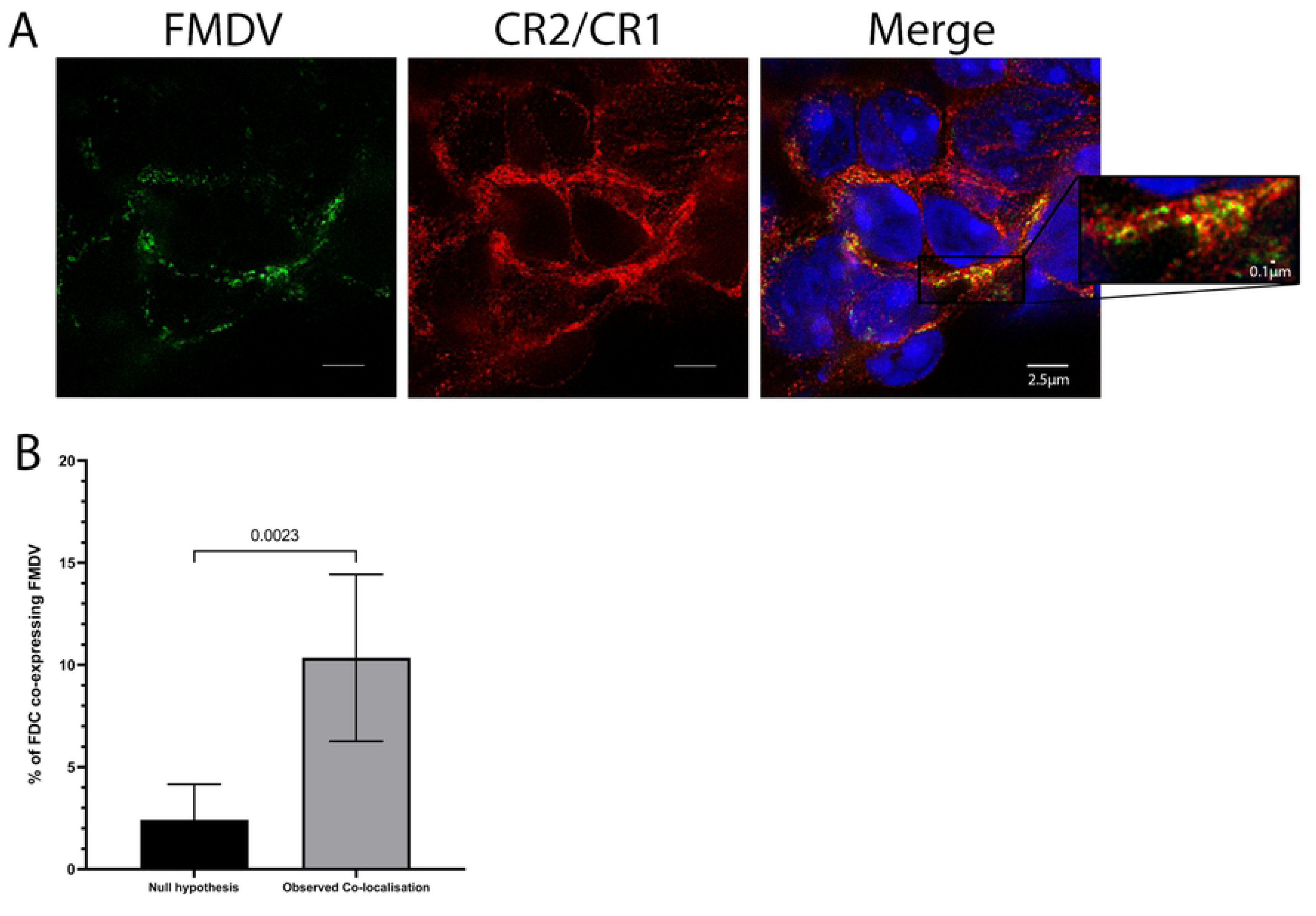
Co-localisation of FMDV with FDCs.

BALB/c mice were infected with FMDV/O/UKG/34/2001 and high-resolution images were taken of spleen samples using a STED confocal microscope. **A**) Spleen taken from an infected mouse 7 dpi, demonstrating the co-localisation of FMDV (green) with FDCs (red). **B**) Morphometric analysis using ImageJ confirmed that FMDV was preferentially associated with FDCs in spleen tissues (n=7) and significantly greater than the null hypothesis that the pixels were randomly distributed, with a p value of 0.0023.

### CR2/CR1-blockade reduces the generation of neutralising Ab in FMDV infected mice

Since the retention of Ag on FDCs is important for the induction and maintenance of high-titre Ab responses and B cell affinity maturation (13, 33), we next tested the hypothesis that CR2/CR1-blockade in FMDV-infected mice would impede the generation of virus neutralizing Ab. Serum samples were collected from mAb 4B2- or control IgG1-treated FMDV-infected mice and incubated with FMDV-susceptible cells and FMDV for their ability to neutralise the virus.

High titres of virus-specific neutralising Ab were detected in the sera of control IgG1-treated mice by 7 dpi, titres increased by day 14 and these were maintained up to 28 dpi (Fig 7). In contrast, while virus-specific neutralising Ab were detected in the sera of mAb 4B2-treated mice by 7 dpi, these did not increase with the infection and their titres were significantly reduced when compared to those in the serum of control IgG1-treated mice (Fig 7).

**Fig 7.**
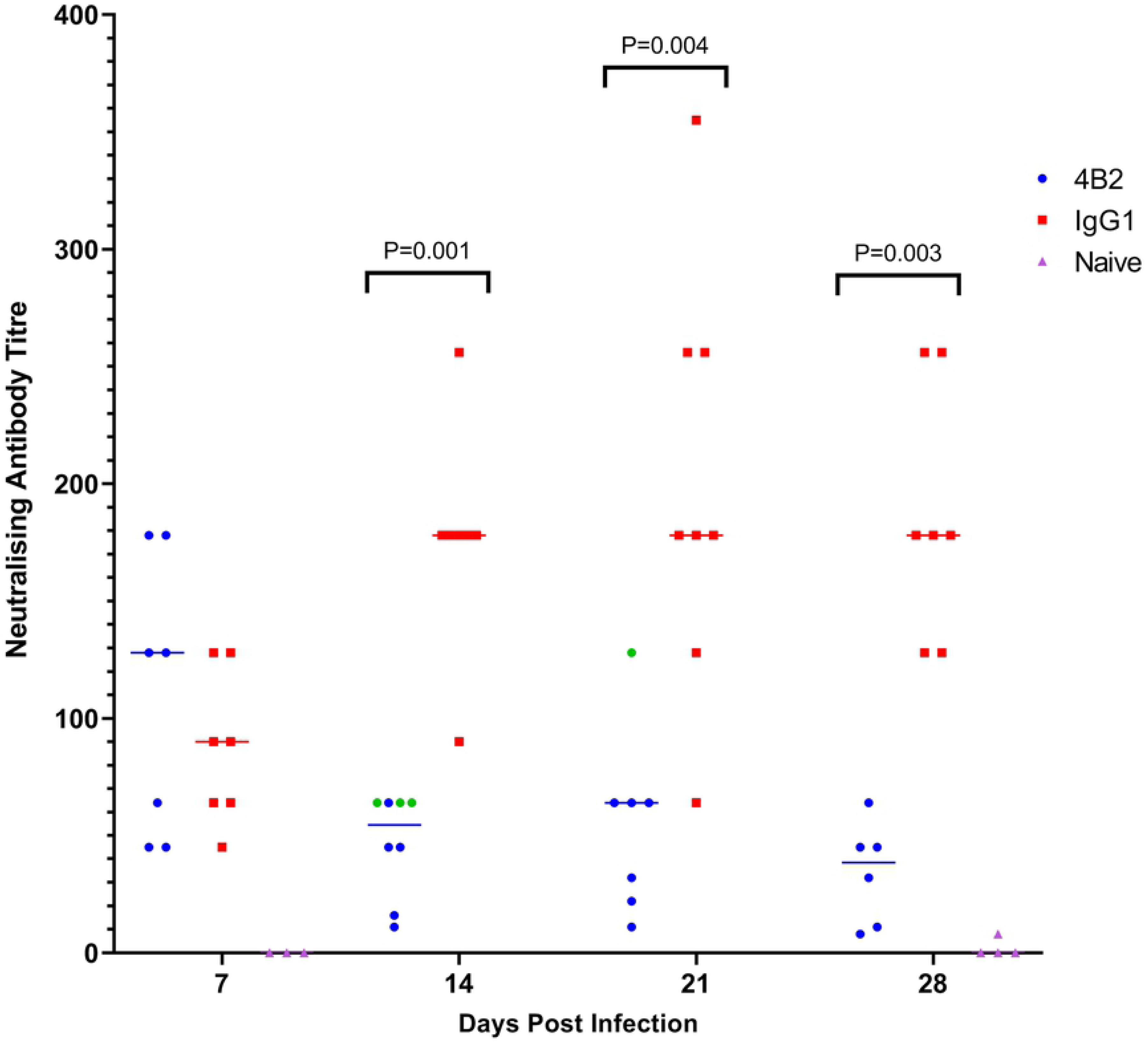
Effect of 4B2 treatment on titres of FMDV neutralising antibodies in mouse serum.

FMDV neutralising antibodies were evaluated from serum samples taken from BALB/c mice at 7, 14, 21 and 28 days post infection with FMDV. Mice had either been pre-treated with mAbs 4B2 or IgG1 1 day prior to FMDV infection. Naïve mice were used as controls. Each blot represents an individual animal and the line represents the median antibody titre.

Neutralising antibody titres are expressed as the serum dilution that neutralised 50% of 100 TCID50 of the virus. The points in green are the mice with FMDV positive GCs from the 4B2 group (Fig 5). Using the non-parametric Mann-Whitney *U* test to compare the medians of the two groups, at 14 dpi p value of 0.001; 21 dpi p value of 0.004 and 28 dpi p value of 0.003.

### CR2/CR1-blockade had no effect on the total IgG/IgM FMDV-specific Ab titres

We next used indirect ELISAs to determine the isotypes of the FMDV-specific Abs produced in the sera of mice from each treatment group. Despite the significant decrease in the level of virus-neutralizing antibodies in the sera of the mAb 4B2-treated mice, there were no significant differences in the titre of virus-specific IgG produced at any of the time points analysed (Fig 8A). An FMDV mAb of known concentration was used in the ELISA as a standard to determine the concentration of FMDV-specific IgG in the polyclonal sera (Fig 8B). At 7 dpi, 4 mice from the 4B2 group and 3 mice from the IgG1 group had FMDV-specific IgM antibodies, and as expected no mice had IgM titres after this timepoint (Fig 8C).

**Fig 8.**
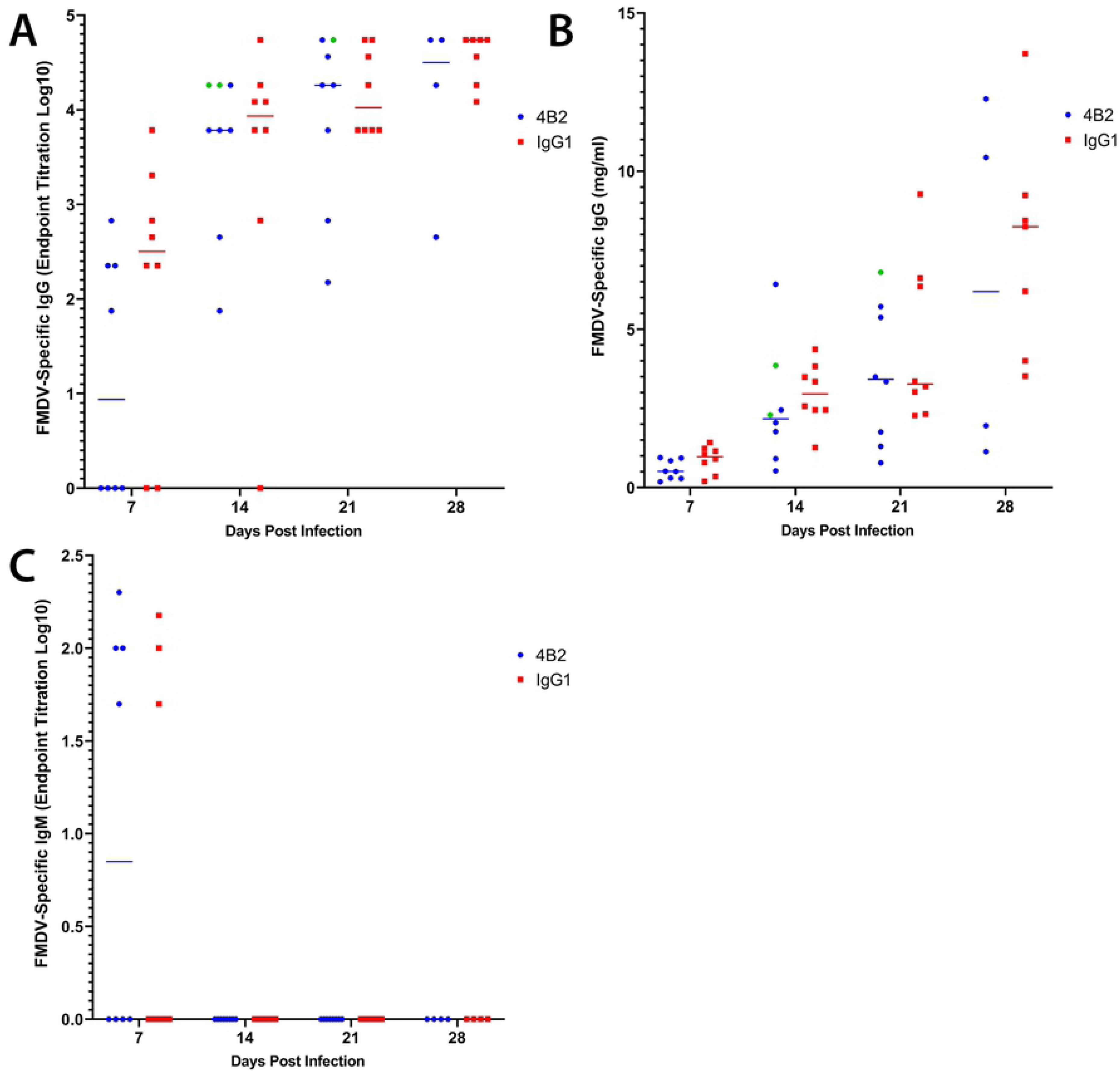
Effect of 4B2 treatment on titres of FMDV specific antibodies.

FMDV-specific antibodies in serum samples from mice treated with 4B2 or an isotype control antibody (IgG1) were detected by ELISA. Serum samples were collected at 7, 14, 21 and 28 dpi and tested for (**A**, **B**) IgG and (**C**) IgM antibodies. Antibody titres were either expressed as (**A**, **C**) the reciprocal log10 of the last positive dilution or (**B**) using a known FMDV IgG standard to plot the concentration of IgG antibodies in mg/ml. Each data point represents an individual animal and the bars represent median values. The points in green are the mice with FMDV positive GCs from the 4B2 group (Fig 5), these mice are included in the statistics. Using the non-parametric Mann-Whitney *U* test to compare the medians of the two groups, there were no statistically significant differences at any time points.

### CR2/CR1-blockade reduces antibody titres to the neutralising FMDV G-H loop

Next, an ELISA was carried out to compare the antibody titres in the mAb 4B2-treatment group and the control group against the O/UKG/12/2001 VP1_129-169_ G-H loop (Fig 9A). In mice and cattle, the G-H loop is a neutralising epitope of FMDV, and a G-H loop peptide vaccine is sufficient to protect mice against FMDV challenge (34, 35). Mice treated with mAb 4B2 had significantly lower antibody titres to the G-H loop compared to the control group, which correlated with the decreased ability of the antibodies from the 4B2-treated mice to neutralise FMDV.

**Fig 9.**
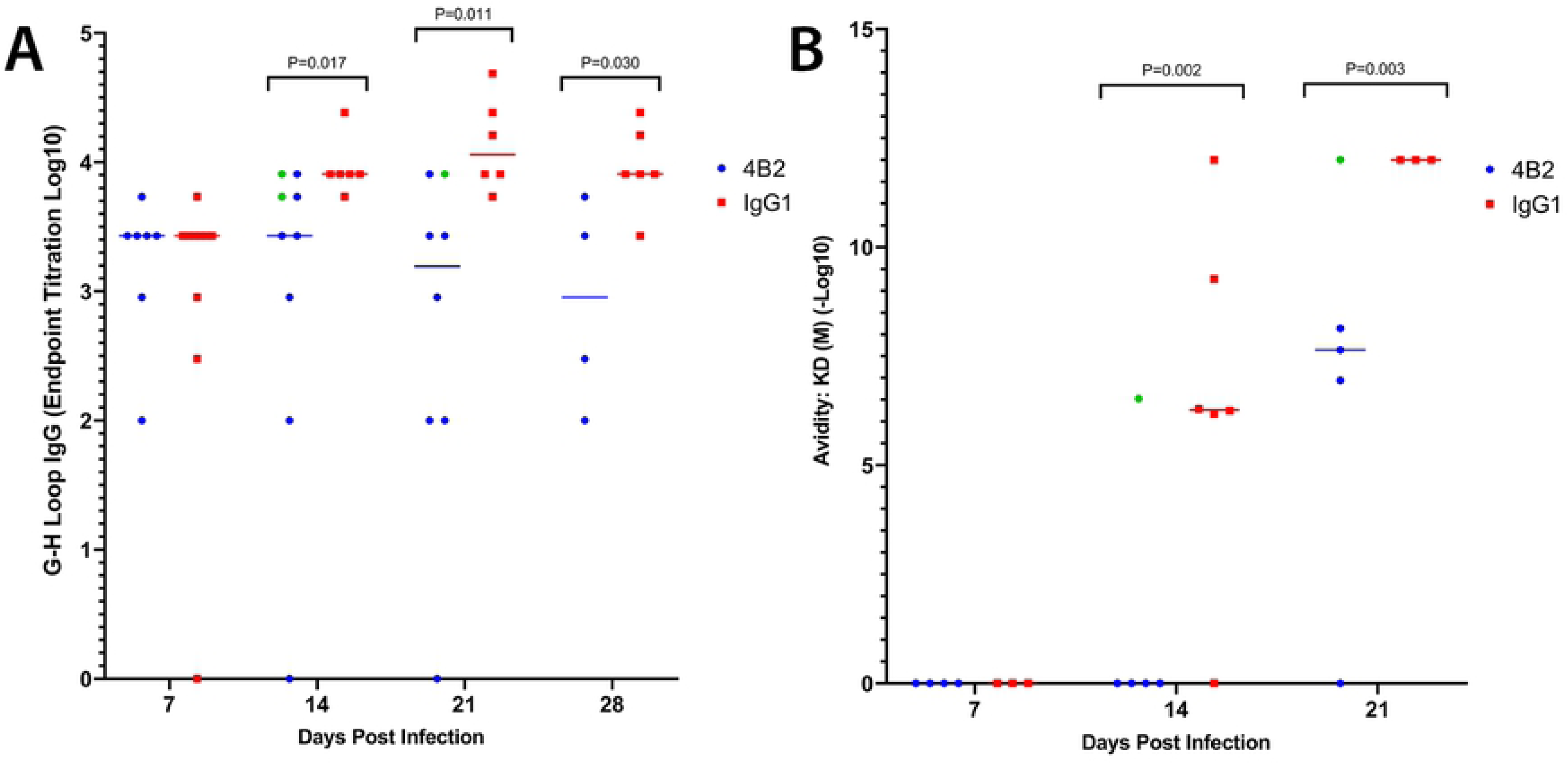
Effect of 4B2 treatment on titres of IgG antibodies specific to the FMDV G-H loop and the avidity of FMDV specific IgG antibodies in mouse serum.

BALB/c mice treated with 500 μg of either 4B2 mAb or IgG1 control mAb on day -1 before challenge with FMDV and sera was collected at 7, 14, 21 and 28 dpi. **A**) An indirect peptide ELISA showed that mice treated with 4B2 had significantly less antibodies to the FMDV G-H loop compared to the IgG1 control group, with a P value of ≤0.05 using the non-parametric Mann-Whitney *U* test. **B**) The avidity of antibodies was measured using biolayer interferometry and was performed using an Octet Red96e. FMD O1/Manisa/TUR/69 VLP were bound to streptavidin sensors and dipped into three dilutions of sera per mouse. Each blot represents the mean avidity of these measurements for each individual mouse, represented as -Log10 of the KD (M) value. Sera which produced a negative response rate, and therefore had too few antibodies bound to FMD VLPs to reach the limit of detection, are recorded as 0. The results demonstrate that mice treated with 4B2 had significantly lower avidity antibodies compared to the control group, with a P value of ≤0.05 using the non-parametric Mann-Whitney *U* test. The points in green are the mice with FMDV positive GCs from the 4B2 group (Fig 5), these mice are included in the statistics.

### CR2/CR1-blockade decreases antibody avidity to FMD VLPs

We then investigated whether CR2/CR1-blockade had affected the avidity of the FMDV-specific Ab for FMDV-Ag (Fig 9B). Using the data shown in Fig 8B, known concentrations of FMDV-specific IgG in polyclonal sera from infected mice from each group were incubated with stable FMD virus-like particles (FMD VLP) and the Ab dissociation/association rates (k_off_/k_on_) rates and *K*_D_ values determined. The *K*_D_ is the equilibrium dissociation constant between an Ab and its Ag and is measured using the ratio of k_off_/k_on_, therefore *K*_D_ values were used to represent the avidity of the polyclonal Abs, based on the individual affinities of the Abs in the polyclonal serum samples, to the FMD VLP. These data clearly showed that the *K*_D_ values in the sera of mice treated with mAb 4B2 were significantly lower than those in the sera of IgG1-treated control mice (Fig 9B); suggesting that virus-specific Ab induced after CR2/CR1-blockade had reduced avidity to FMD VLP.

## Discussion

Our previous studies suggested that in cattle FMDV is localised on FDCs in the light zone of GCs (16). Similar to studies with HIV where the interaction of virus with FDCs has been explored in detail in mice, we have demonstrated FMDV localises to FDCs in mice after the resolution of viraemia. The mouse model for FMDV persistence showed FMDV Ag in the GC for up to 63 dpi, associated with FDCs (36). We have now used this FMDV mouse model to gain novel insight into the mechanisms of FMDV persistence on FDCs. In this study, FMDV protein was detected in GC up to 28 dpi and FMDV genome up to 21 days in spleen samples. We suspect the absence of detectable genome at 28 days is because the RNA will be in a small number of localised deposits in the GC which may not be detected when the whole spleen is sampled. We have shown previously in cattle and African buffalo that FMDV genome does persist in GC for prolonged periods (16, 25).

The absence of FMDV Ag and genome by IHC and PCR in mice treated with mAb 4B2 as reported here, demonstrates the role of CR2/CR1 as the major receptor involved in the trapping and retention of FMDV. Furthermore, blocking of CR2/CR1 results in a significant reduction of neutralising antibody titres against FMDV. Although two mechanisms have been described for antigen trapping by FDC, CR mediated (14) and FcR mediated (37), the near complete elimination of FMDV on FDCs after treatment with 4B2 leads us to believe the trapping is CR2/CR1-dependent. However, we do not exclude that longer persistence of the virus on FDCs after natural infections, when anti-virus antibody forms, are also due to FcR accompanying CR2/CR1.

A similar study by Gustavsson et al. found that blocking CR2/CR1 led to an inhibition of both the primary Ab response and the induction of memory B cells to a T-I Ag. Immune complexes bound to FDCs via CR2/CR1 were required for efficient T-I B cell stimulation (38). Ochsenbein et al. used Cr2^-/-^ mice to investigate antibody responses to a T-I Ag, VSV. They showed similar findings, that early antibody responses to infection were unaffected in these knockout mice, including no significant effect on the IgM response to infection in mice deficient in CR2/CR1. However, longer term antibody responses to VSV were not significantly different in Cr2^-/-^ mice compared to the wild type (WT) (39). Unlike FMDV (12), VSV is able to induce B cell memory, therefore, the contrast to our findings could be because the induction of antibody responses to VSV are less dependent on antigen persistence on FDCs compared to FMDV. The murine Cr2 gene encodes two proteins, CR1 and CR2, via alternative splicing (40), therefore inactivation of the Cr2 gene leads to deficiency in both CR1 and CR2. The similarities in these receptors also leads to blocking of both CR1 and CR2 upon administration of an anti-CR2 and/or -CR1 mAb.

Cr2^-/-^ mice have abnormalities in the maturation of GCs including the GC B cells associated with the CR2/CR1 deficiency, which may complicate the interpretation of some studies where they are used. These mice have been shown in multiple studies to have a discernible impairment in their ability to mount a humoral immune response (41, 42). A recent study by Anania et al. used image analysis to demonstrate that FDCs lacking CR1 and CR2 not only have a decreased ability to capture ICs, but in the Cr2^-/-^ mice, GCs are fewer and smaller and FDCs are poorly organised (43). FDCs use cytokine gradients to interact with B cells and T follicular helper cells in GC, therefore disorganisation of the FDC networks leads to a variety of abnormalities, including impaired B cell survival and reduced Ig production (44).

Although Cr2^-/-^ mice are unable to mount a normal humoral immune response to various antigens, a study showed that Cr2^-/-^ mice had reportedly normal levels of total IgM and of the different IgG isotypes, showing no evidence of altered B- or T- cell development (42). These studies showed antibody titres were similar in Cr2^-/-^ and wildtype (WT) mice, however functional differences in antibodies were not specifically investigated. We also showed IgM and IgG titres were similar in treated and control mice, but went on to show low avidity, non-neutralising antibodies were produced which could be due to a defective affinity maturation process due to the lack of binding of FMDV proteins to CR2/CR1 on FDCs. Furthermore, it has been established that in mice the G-H loop is a neutralising epitope inducing protection against FMDV (34); and mice treated with 4B2 had lower titres of antibodies to the G-H loop. These results correlate with the reduced ability of the antibodies from the mice treated with 4B2 to neutralise FMDV from 7 dpi.

Due to the off-target effects from using Cr2^-/-^ knockout mice to study the function of FMDV antigen on FDC, we used the 4B2 monoclonal antibody to block CR2/CR1 on FDCs. This antibody had been previously described to block these receptors for up to 6 weeks *in vivo* in mice, without disrupting other cell types. Bioimaging and flow cytometry analysis confirmed that GC numbers and sizes were normal and the percentage of other immune cell subsets in the spleen were unaltered after blocking up to 35 days, including B- and T- cells. We were therefore confident that the 4B2 mAb would indicate whether antigen bound to CR2/CR1 on FDCs impacted on the immune response.

A number of studies have used *Cr2^-/-^* mice and reconstituted with *Cr2^+/+^* WT bone marrow (BM) to allow a more specific investigation of the role of CR2/CR1 on FDCs without impairing B cell functions (45–47). This is possible because FDCs are derived from stromal cells; whereas B cells are BM in origin. Initial IgG and IgM responses were shown to be similar in *Cr2^-/-^* mice with or without WT BM (Cr2^+/+^ B cells), suggesting Ag can induce a B cell response in the absence of CR expression (45, 46). However, studies investigating the long-term antibody response of these chimeric mice have shown a significant reduction in both long-term antibody production and memory when FDCs specifically did not express Cr2 (46, 47). This is in line with our results where neutralising antibody responses up to day 7 post infection were similar in mice with or without a CR2/CR1 block, yet after this timepoint, there was a significant reduction in FMDV neutralising antibodies in mice treated with the anti-CR2/CR1 mAb.

It is well established that CR1 and CR2 are essential for binding ICs and are expressed at high levels on FDCs; and while FDCs can also trap ICs via the FcR, it is to a lesser degree [14-16, 48]. FDCs can acquire antigen through various pathways, including direct interaction by small antigens as well as by binding to complement component 3 (C3) fragments on ICs via CR2/CR1 when presented to them via B cells (17, 48). It has been previously described that C3 fragments, specifically C3d, could therefore be used as a vaccine adjuvant (49). A study by Ross et al. demonstrated the effectiveness of C3d-fusions to haemagglutinin in enhancing antibody production and maturation, leading to a protective immune response in the influenza mouse model (50). This would be a particularly interesting area of research for FMDV, due to the short duration of immunity after FMD vaccination. If fusion of C3d to FMD vaccine antigens resulted in targeted antigen deposition on FDCs, this could improve the magnitude and duration of the neutralising antibody response.

We have been unable to demonstrate that FMDV retained by FDCs is infectious. Heesters et al. found that FDCs retained infectious virus in cycling endosomes and this compartmentalisation of virus could be the reason for our observations (23). Ultrastructural studies are planned to establish whether intact virus is present on FDCs in the splenic GCs and to determine whether under certain circumstances FMDV on FDCs is able to infect cells, or whether the virus becomes attenuated on FDCs. This will not only provide insight into the

mechanisms of persistence, but whether virus immune-complexed onto FDCs is infectious and therefore whether there is a possibility of re-infection or transmission of virus. This knowledge could therefore potentially be useful for future decision making for controlling FMDV outbreaks.

## Materials and Methods

### Mice and experiment design

Experiments were carried out to address 3 objectives; firstly to determine whether the 4B2 mAb successfully blocked CR2/CR1 by using PAP which is known to bind to FDCs via the CR2/CR1. Secondly, to determine the effect of 4B2 on the cell subsets of the spleen.

Finally, a challenge study to determine whether FMDV needs to bind to FDCs via the CR2/CR1 to maintain a neutralising antibody response. Female BALB/c mice (8-12 weeks) were used in these experiments and were purchased from Charles River Laboratories, UK. Mice were acclimatised for 7 days before being used in experiments and were maintained with food and water ad-libitum and full environmental enrichment. Mice were humanely culled using isoflurane and a rising concentration of carbon dioxide (CO_2_) method. All animal experiments were performed in the animal isolation facilities at the Pirbright institute and were conducted in compliance with the Home Office Animals (scientific procedures) ACT 1986 and the Pirbright Institute’s Animal Welfare and Ethical review procedure.

#### 4B2 treatment

BALB/c mice were given a single intraperitoneal (i.p.) injection of 200µl of 0.5 mg purified mAb 4B2 to mouse CR2/CR1 (30). Animals treated with the same dose of a mAb anti-OVA IgG1, F2.3.58 antibody (2B Scientific, UK) were used as isotype matched controls. Two mice treated with 4B2 and two with the IgG1 control mAb were culled at 2 and 7 days post treatment “early time points”, and a further two from each group at 22 and 35 days post treatment “late time points” to assess the effects of 4B2 on spleen cell subsets. The spleen samples were collected in RPMI media (Gibco, UK) and immediately processed in the lab for flow cytometry.

#### PAP treatment

To test the ability of 4B2 to block CR2/CR1 *in vivo*, mice were given a single injection of 100µl preformed rabbit peroxidase-anti-peroxidase (PAP) immune complexes (Sigma) intravenously (i.v.) 1 day after treatment with 4B2 (n=4) or anti-OVA IgG1 (n=4). Mice were culled 1 day later and their spleens were collected in optimal cutting temperature (OCT) compound (VWR Chemicals, UK) and stored at -80° C to test for the presence of FDC-associated IC by confocal microscopy.

#### FMDV infection

1 day after treatment with 4B2 or IgG1 mAbs, mice were inoculated i.p. with a total dose of 10^6.2^ TCID_50_ of FMDV/O/UKG/34/2001 in 200µl. After challenge, mice were bled from the tail vein at 2 dpi. Terminal bleeding after culling from cardiac puncture and spleens from culled mice were collected from 8 animals at 7, 14, 21 and 28 dpi from each treatment group. The spleens were cut in half, with half collected in OCT for analysis by confocal microscopy and half collected in RPMI medium (Gibco, UK) for analysis by PCR. The whole blood samples were stored at 4° C overnight to allow blood clotting, the samples were centrifuged, and the serum was collected and stored at -80° C.

### Processing splenocytes

The spleen samples collected in RPMI medium were homogenised and passed through 70 µm cell mesh strainers (BD Biosciences, UK). Cells were washed in RPMI medium by centrifugation and red blood cells were lysed with ACK lysing buffer (Sigma-Aldrich, UK). Following lysis, cells were washed twice in RPMI by centrifugation and re-suspended in RPMI complete medium (RPMI with 10% foetal bovine serum (Gibco, UK), 1% Gibco penicillin-streptomycin (10,000 U/ml) (Life Technologies, UK) and 1% Gibco MEM non-essential amino acids (100X) (Life Technologies, UK)), counted and stored at 4° C overnight prior to flow cytometric analysis.

### Flow Cytometry

The processed splenocytes were distributed at 1 x 10^6^ per well into Nunc 96-well round bottom microwell plates (Thermo Scientific, UK). The cells were blocked by adding 5µg/ml purified rat anti-mouse CD16/CD32 (mouse BD Fc Block) clone: 2.4G2 (BD Biosciences, UK) in autoMACS buffer (Miltenyi Biotec, UK). Cells were stained with CD8a-FITC (Life Technologies), CD4-PE (Miltenyi Biotec) to detect cytotoxic and helper T cells respectively and B220 biotin RA3-6B2 Alexa Fluor 647 to detect B cells (CD45R), CD11b-APC to detect dendritic cells and CD169 (Siglec-1)-APC to detect marginal zone macrophages. Streptavidin Molecular Probe Alexa-Fluor-633 conjugated secondary mAb (1µg/ml) (Invitrogen) was used to detect biotinylated antibodies 7E9 (BioLegend, UK) and 7G6 (BD Biosciences, UK) to identify CD21/CD35 (CR2/CR1) and 8C12 (BD Biosciences, UK) to identify CD35 (CR1). Single staining controls and no staining controls were also included for compensation purposes. The cells were then fixed with 1% paraformaldehyde, washed and resuspended in MACS buffer, before being read on the MACS Quant (Miltenyi Biotec, UK). The analysis was completed using FCS Express (De Novo Software, US).

### Quantification of viraemia by plaque assay

Foetal goat tongue cells (ZZR cells), which are highly susceptible to FMDV, were grown up to 95-100% confluency in 6 well plates. Cells were washed in PBS and a 10-fold dilution of serum samples from 2 dpi (n=30 and n=22 from 4B2 and IgG1 treatment group, respectively) and 7 dpi (n=4 per treatment group) were added to the wells. Serum from 2 naïve animals at 2 dpi and 1 naïve animal at 7 dpi were used as negative controls. Plates were incubated for 30 minutes at 37℃ with 5% CO_2_ and then 3ml/well of Eagle’s Overlay-Agarose (Eagle’s overlay media (TPI, UK) and 2% agarose (Sigma, UK)) was added and allowed to set at room temperature. Plates were incubated at 37℃ with 5% CO_2_ for 48 hours. Following incubation, plates were fixed and plaques visualized by staining the cell monolayer with methylene blue in 4% formaldehyde in PBS for 24 hours at room temperature. The plates were washed with water and the agarose plugs discarded. The viraemia was expressed as the Log10 of the number of plaque forming units per ml (PFU/ml).

### One Step RT-qPCR of serum samples

Due to low volumes of serum collected from the tail vein, serum samples from animals taken at 2 dpi were pooled to reach 50µl. Therefore, FMDV genome copy number was measured by RT-qPCR in 6 pools of serum from IgG1 and 4B2 treated mice and 3 pools from the naïve groups. Fifty µl of serum samples taken from terminal bleeds from culled animals at 7 dpi were also analysed. The RNA was extracted using the MagVet^TM^ Universal Isolation Kit (Thermo Fisher Scientific, UK) and the KingFisher^TM^ Flex (Thermo Fisher Scientific, UK). The PCR was performed, using the SuperScript™ III Platinum™ One-Step Callahan 3D quantitative RT- qPCR, according to the standard protocol of the World Referenced Laboratory for FMDV, with a cut-off cycle threshold (Ct) of ≥ 35 (51). Results were expressed as Log10 FMDV genome copy number (GCN)/ml of sample by extrapolating the Ct values to GCN by using a linear regression model with serial dilutions of in vitro synthetized 3D RNA standard.

### RT-qPCR from tissues

Spleen samples were homogenised in 200μl DMEM media (Gibco, UK) using the FastPrep-24 and lysing matrix tubes (MP Biomedicals) prior to RNA extraction (as described above). Following RNA extraction, cDNA was generated using TaqMan reverse transcription reagents (Applied Biosystems, UK,). The EXPRESS qPCR SuperMix Universal Kit (Invitrogen, UK) was used for real time-PCR and the PCR reactions for FMDV 3D were performed as previously described, with a cut-off cycle threshold (CT) of ≥ 35 (51). The 18S ribosomal RNA housekeeping gene was used for normalisation based on previously published primers (52, 53). The PCR reaction was performed on a Stratagene MX3005p quantitative PCR instrument (Stratagene, USA). Results were expressed as Log10 FMDV RNA copies/10^8^ copies 18S.

### Immunofluorescence by confocal microscopy

The frozen spleens embedded in OCT were cut on a cryostat (7-9 μm), mounted on a superfrost slide and stored at -20° C overnight. The slides were air-dried, fixed with 4% paraformaldehyde and blocked with 5% normal goat serum (NGS) (abcam, UK). FDC networks visualised by staining with 1µg/ml Alexa Fluor 594-conjugated anti-mouse CD21/CD35 (CR2/CR1) antibody, clone 7E9 (BioLegend, UK), marginal zone macrophages were visualised using 1µg/ml CD169 (Siglec-1), clone MOMA-1 (Bio-Rad, UK), 2µg/ml biotinylated llama single domain anti-FMDV 12S antibody VHH-M3 (Kindly provided by Dr M Harmsen, Central Veterinary Institute of Wageningen, AB Lelystad, The Netherlands) (54) was used to detect FMDV/O/UKG/34/2001 and goat anti-rabbit Molecular Probe Alexa-Fluor-488 was used to detect PAP IC. Goat anti-rat and streptavidin Molecular Probes Alexa-Fluor-488 and 633 conjugated secondary mAbs (Invitrogen) were used at 2µg/ml and all sections were counterstained with DAPI to distinguish cell nuclei. Spleen sections were visualised, imaged and all data was collected using a Leica SP8 confocal microscope (Leica Microsystems GmbH, Germany).

The same protocol was used for the stimulated emission depletion (STED) with the following changes: goat anti-rat and streptavidin Molecular Probes Alexa-Fluor-488 and 555 conjugated secondary mAbs (Invitrogen) were used at 4µg/ml, ToPro3 was used for nuclear staining and a super-resolution Leica TCS SP8 STED 3X microscope (Leica Microsystems GmbH, Germany) equipped with 592 and 660nm depletion lasers was used to image and collect data. STED images were then deconvolved in Huygens Professional software 21.04 (Scientific Volume Imaging, Netherlands) using the Deconvolution Wizard with a theoretical PSF. Data was analysed using ImageJ software as previously described (31, 32) to compare the null hypothesis (that the pixels were randomly distributed) to the observed levels of co-localisation.

### Virus Neutralising Test

Serum samples collected at 7, 14, 21 and 28 dpi were heated at 56° C for 1 hour to inactivate complement and analysed for their ability to neutralise a fixed dose of FMDV on IB-RS-2 cells porcine cells). Samples were then diluted 2-fold in 96 well plates in duplicate in serum free medium starting from a 1:8 dilution. Naïve mouse serum and cells only were used as negative controls. One hundred tissue culture infectious dose 50 (TCID_50_) of FMDV OUKG was added to all wells excluding cell only controls. Plates were incubated for 1 hour at room temperature before 5 × 10^4^ IB-RS-2 cells were dispensed to each well. The plates were incubated at 37°C in 5% CO2 and checked daily for cytopathic effect (CPE). After 72 hours the plates were inactivated with 1% Trichloroacetic acid (TCA) (Sigma-Aldrich, UK) washed with water and stained with methylene blue. Neutralising antibody titre was calculated using the Spearmann-Karber formula and results expressed as the log_10_ reciprocal serum dilution that neutralised 50% of 100 TCID50 of the virus (55).

### IgG and IgM ELISA

An indirect enzyme-linked immunosorbent assay (ELISA) was developed to detect FMDV-specific mouse antibodies. The assay was adapted from the FMDV isotype specific ELISA protocol to detect antibodies to FMDV in cattle and swine serum (10). ELISA plates were coated with a rabbit anti-O FMDV polyclonal antibody (TPI, UK), washed with PBS containing 0.05% Tween20 (Sigma, UK) and then 0.5 μg/ml inactivated FMDV/O1 Manisa vaccine antigen (Merial, UK), diluted in blocking buffer (1:1 PBS and SEA BLOCK (Thermo Scientific, UK)), was added to each well. Serum samples were added, and bound antibodies were detected by incubating the plates with horseradish peroxidase-conjugated goat anti-mouse IgG or IgM (Invitrogen, UK), diluted in blocking buffer. TMB substrate (Thermo Scientific, UK) was used as a developer and the reaction was stopped with 0.3M H_2_SO_4_ and the optical density (OD) was read at 450 nm. Antibody titres were expressed as either log10 of the reciprocal of the last dilution with a mean OD greater than 1.5 times the mean of the OD of the negative control serum or using an FMDV-specific IgG standard (IB11 mAb (10)) of known concentration, a standard curve was generated to determine the concentration of the FMDV-specific IgG in the serum samples analysed. The O1/Manisa/TUR/69 vaccine Ag was used due to good cross-reactivity and cross-protection with OUKG as demonstrated in previous studies (56–58).

### Peptide ELISA

An indirect peptide ELISA using a biotinylated O/UKG/12/2001 G-H loop peptide (VYNGNCKYGESPVTNVRGDLQVLAQKAARTLPTSFNYGAIK) (Peptide Protein Research Ltd, UK) was developed to determine the presence of antibodies directed against the FMDV VP1_129-169_ G-H loop. The highest concentration of each serum sample was also tested with a biotinylated negative control peptide with a similar molecular weight and number of charged residues vs hydrophobic residues (PSRDYSHYYTTIQDLRDKILGATIENSRIVLQIDNARLA) (Peptide Protein Research Ltd, UK) to ensure the sera wasn’t binding non-specifically (data not shown). Streptavidin coated ELISA plates (Thermo Scientific) were incubated with 8 μg/ml G-H loop peptide diluted in PBS at 37°C for 2 hours. The plates were washed with TBS containing 0.1% BSA 0.05% Tween20 (Sigma, UK) and then serum samples were added in duplicate. Bound antibodies were detected by horseradish peroxidase-conjugated goat anti-mouse IgG (Invitrogen, UK) and SIGMAFAST™ OPD (o-Phenylenediamine dihydrochloride) (Sigma, UK). The optical densities (OD) were measured at 450nm and antibody titres were expressed as log10 of the reciprocal of the last dilution with a mean OD greater than 1.5 times the mean of the OD of the negative control serum.

### Biolayer Interferometry

Biolayer interferometry was performed using an Octet Red96e instrument (ForteBio, Inc.) and ForteBio Data Analysis HT software (v 11.1.0.25) was used to determine the response rate, k_off_/k_on_ rates and the *K*_D_ (M) values. This method was adapted from previously described methods using polyclonal sera (59, 60). A 5 μg/ml concentration of biotinylated stable O1/Manisa/TUR/69 FMD VLP (61) (kindly provided by Alison Burman) was immobilised on streptavidin-coated biosensors (Sartorius UK Limited) for 900 s. A baseline was established by measurements taken when sensors were immersed for 60 s in HEPES 10 mM, NaCl 150 mM, EDTA 3 mM, 0.005% Tween 20 (HBS-EP) buffer (Teknova). The sensors were then immersed in a dilution series of polyclonal sera, with known FMDV-specific IgG concentrations, from mice taken at 7, 14 or 21 dpi with FMDV for 1200 s in the association phase. Subsequently, the sensors were immersed in HBS-EP buffer for 1200 s in the dissociation phase. Unloaded sensors and reference wells were used to subtract non-specific binding. Mean *K*_D_ (M) values were obtained from the dilution series of each mouse based on their global fit to a bivalent model, with a full R^2^ value of ≥0.9. The *K*_D_ values were measured using the ratio of k_off_/k_on_, to determine the avidity of antibodies in the polyclonal serum samples to the FMD VLP. The values were expressed as -Log10 of *K*_D_ (M) values, and sera which had a response rate below 0 were recorded as 0.

### Statistical analysis

The comparisons between the experimental groups and their corresponding control groups were carried out using Minitab software (Minitab, US). The non-parametric Mann-Whitney *U* test was used to compare the medians of viremia, presence of antigen, antibody titres and avidities and splenic cell subsets between the 4B2 and IgG1 treated groups. A *P* value of ≤ 0.05 was considered statistically significant.

## Acknowledgments

We thank The Pirbright Institute animal services team, in particular David Selby, for their help with the *in vivo* procedures; Andrew Shaw and Holly Everest for their guidance with the Octet; Barry Bradford for his help with ImageJ; the bioimaging team, namely Jennifer Simpson and the flow cytometry facility.

## Author Contributions

Conceived and designed the experiments: LG EP NM BC. Performed and analysed the data: LG EP JW. Contributed reagents/materials/analysis tools: LG EP BC LK JW. Wrote the paper: LG. Revised the draft for important intellectual content: BC EP NJ NM JW LK. Approved the final version for publication: LG BC EP NM NJ JW LK.

